# The reference genome of the paradise fish (*Macropodus opercularis*)

**DOI:** 10.1101/2023.08.10.552018

**Authors:** Erika Fodor, Javan Okendo, Nóra Szabó, Kata Szabó, Dávid Czimer, Anita Tarján-Rácz, Ildikó Szeverényi, Bi Wei Low, Jia Huan Liew, Sergey Koren, Arang Rhie, László Orbán, Ádám Miklósi, Máté Varga, Shawn M. Burgess

## Abstract

Over the decades, a small number of model species, each representative of a larger taxa, have dominated the field of biological research. Amongst fishes, zebrafish (*Danio rerio*) has gained popularity over most other species and while their value as a model is well documented, their usefulness is limited in certain fields of research such as behavior. By embracing other, less conventional experimental organisms, opportunities arise to gain broader insights into evolution and development, as well as studying behavioral aspects not available in current popular model systems. The anabantoid paradise fish (*Macropodus opercularis*), an “air-breather” species from Southeast Asia, has a highly complex behavioral repertoire and has been the subject of many ethological investigations, but lacks genomic resources. Here we report the reference genome assembly of *Macropodus opercularis* using long-read sequences at 150-fold coverage. The final assembly consisted of ≈483 Mb on 152 contigs. Within the assembled genome we identified and annotated 20,157 protein coding genes and assigned ≈90% of them to orthogroups. Completeness analysis showed that 98.5% of the Actinopterygii core gene set (ODB10) was present as a complete ortholog in our reference genome with a further 1.2 % being present in a fragmented form. Additionally, we cloned multiple genes important during early development and using newly developed *in situ* hybridization protocols, we showed that they have conserved expression patterns.

## Background

During the 20^th^ century experimental biology gained increased influence over descriptive biology and concomitantly most research efforts began to narrow into a small number of “model” species. These organisms were not only selected because they were considered to be representative models for the examined phenomena but were also easy and cheap to maintain in laboratory conditions [1, 2]. Working with these convenient experimental models had several advantages and made a rapid accumulation of knowledge possible. It enabled scientists to compare and build on each other’s findings efficiently as well as to share valuable data and resources that accelerated discovery. As a result of this, a handful of model species have dominated the field of biomedical studies.

Despite their broad success, these models also brought limitations. As Bolker pointed out: “The extraordinary resolving power of core models comes with the same trade-off as a high-magnification lens: a much-reduced field of view” [3]. In the case of zebrafish research this trade-off has been perhaps most apparent for behavioral studies. Zebrafish are an inherently social (shoaling) species, but most behavioral studies use them in solitary settings, which arguably is a non-natural environment for them. Therefore, the use of other teleost species with more solitary behavioral profiles is warranted for studies of individual behaviors.

Paradise fish (*Macropodus opercularis* Linnaeus, 1758) is a relatively small (8-11 cm long) freshwater fish native to East Asia, where it is commonly found in shallow waters with dense vegetation and reduced dissolved oxygen [4]. Similar to all other members of the suborder *Anabantoidae*, they are characterized by the capacity to take up oxygen directly from the air through a highly vascularized structure covered with respiratory epithelium, the labyrinth organ (LO) [5]. The ability to ”air-breathe” allows anabantoids to inhabit swamps and small ponds with low levels of dissolved oxygen that would be impossible for other fish species, therefore the LO can be considered an adaptation to hypoxic conditions [6]. The evolution of the LO has also improved hearing in some species [7, 8], and may have led to the emergence of novel and elaborate mating behaviors, including courtship, territorial display, and parental care [6, 9]. The intricate and complex behavior of these fish made them an important ethological model during the 1970-80s, which resulted in a detailed ethogram of the species [10, 11].

We propose that with recently developed husbandry protocols [12] and the advent of novel molecular techniques for genome editing and transgenesis, paradise fish could become an important complementary model species for neurogenetic studies [13]. Furthermore, several genomes are now available for the Siamese fighting fish (*Betta splendens*), a sister species of paradise fish [14–16], so a good quality genome sequence of paradise fish would enable comparative ecological and evolutionary (eco-evo) studies.

While the mitochondrial genome was already available for this species [17] a full genome sequence was lacking. Here, we provide a brief description and characterization of a high quality, *de novo* paradise fish reference genome and transcriptome assembly. We also validate our findings by establishing protocols to perform *in situ* hybridization experiments to detect the spatio-temporal expression of various transcripts and subsequently describing the gene expression patterns for several paradise fish orthologs of genes important in zebrafish development.

## Results and discussion

### Assembly quality and completeness

We generated the *de novo* reference genome sequence for this species using 150X coverage of PacBio SMRT HiFi long-read sequencing and the HiFiasm genome assembly pipeline [18]. The final assembly consisted of 483,077,705 base pairs (bp) on 152 contigs (Supplementary Table, S1). The assembled genome demonstrated a very high contiguity with an N50 of 19.2 megabases (Mb) in 12 contigs. The largest contig was 24,022,457 bp and the shortest contig was 14,205 bp. More than 98% of the canonical k-mers were 1x copy number indicating that our genome assembly is of very good quality (Figure 1A). The paradise fish genome repeat content is estimated to be ∼10.4%. The “trio binning” [19] mode of HiFiasm was attempted using single nucleotide variant (SNV) data collected from short read sequencing of the F_0_ parents, however the heterozygosity rate from the lab raised fish was very low at ∼0.07% making it impossible to efficiently separate maternal and paternal haplotypes. The resulting assembled reference genome is therefore a pseudohaplotype. The sequence of the mitochondrial genome (mtDNA) was essentially identical to the previously published mtDNA sequence for this species (16,495/16,496 identities) [17]. We performed a whole genome alignment to a recent *Betta splendens* assembly [20] and as expected most chromosomes had a 1 to 1 relationship with the exception of *B. splendens* chromosomes 4 and 9 aligning to two separate paradise fish contigs (Figure 1B). This is explained by the number of chromosomes for each species with *B. splendens* having 21 and *M. opercularis* reportedly having 23 chromosomes [21]. We have not determined whether the *B. splendens* chromosomes fused or if the *M. opercularis* chromosomes split.

**Figure 1:**
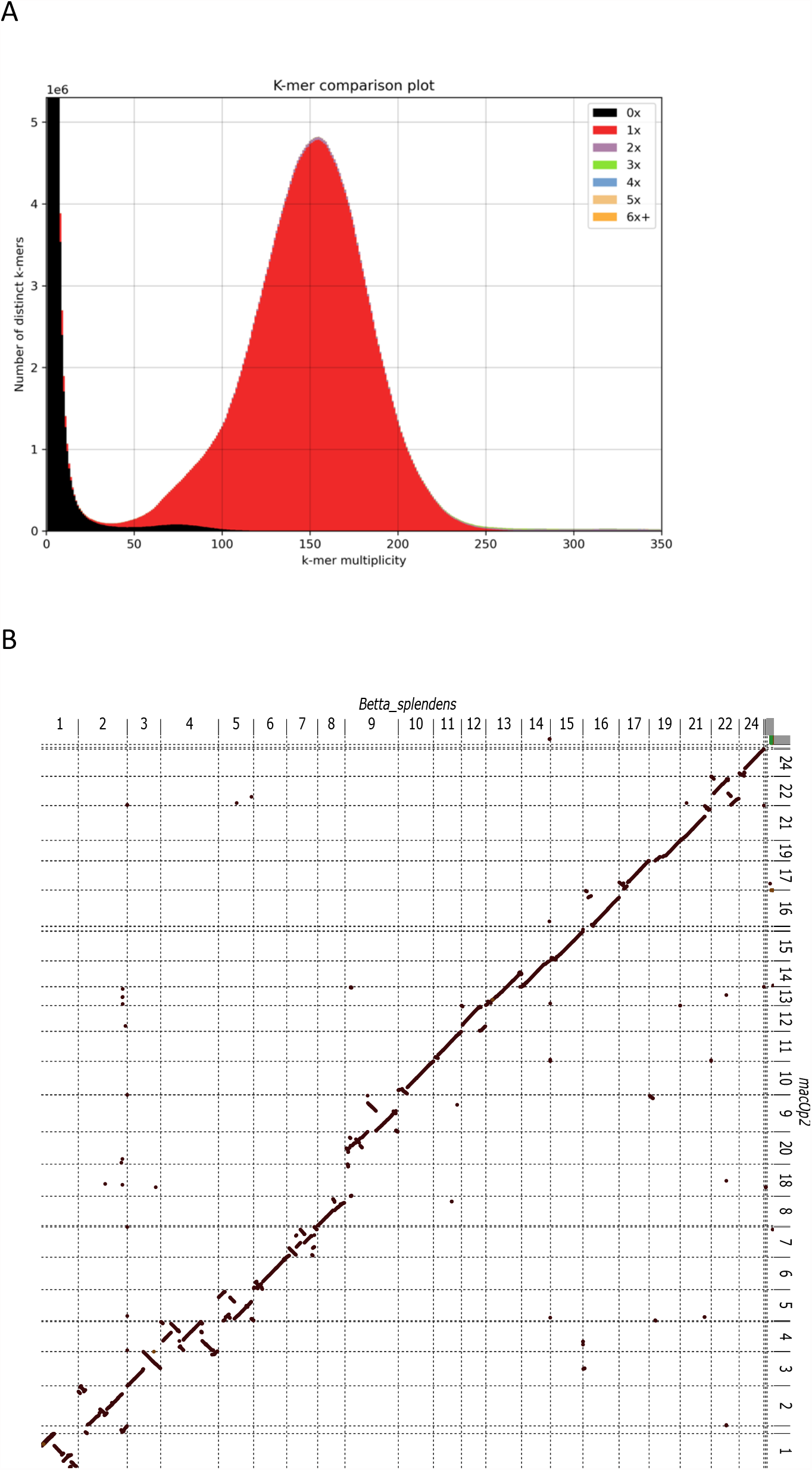
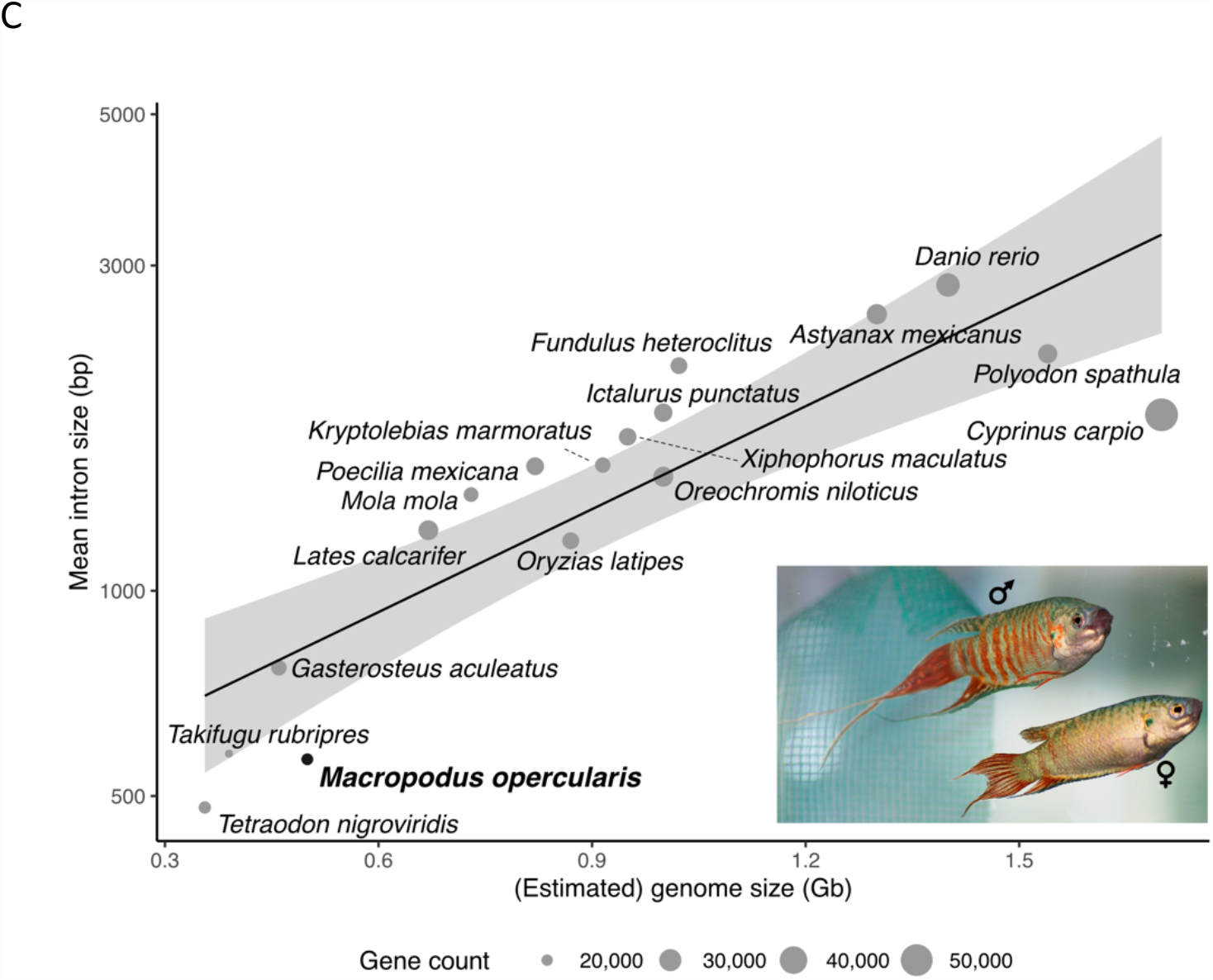
Basic genome assessment. A) K-mer comparison plot. It shows copy number of the k-mers as a stacked histogram colored by the copy numbers found in the paradise fish draft genome assembly. The y-axis shows the number of distinct k-mers, and the x-axis shows the k-mer multiplicity (coverage). Most k-mers are represented once (red peak) indicating high quality for the genomic assembly. B) Whole genome alignment of the *B. splendens* genome assembly to the *M. opercularis* assembly. *B. splendens* is on the X-axis and *M. opercularis* on the Y-axis. *B. splendens* chromosomes 4 and 9 appear to map to two separate *M. opercularis* chromosomes. Chromosome numbering is based on *B. splendens* alignment to medaka chromosomes [14,20]. C) Distribution of intron sizes for various species of fish showing that *M. opercularis* is at the extreme low end of the spectrum, similar to puffer fish species. The line denotes the linear regression line fitted over the data. Grey areas denote the 0.95 confidence interval of the fit. The diameter of the circles correlates to the approximate gene count (20 to 50 thousand).

The genome is relatively compressed in size compared to those of most sequenced teleosts. As expected, the small size of the paradise fish genome is partly due to the relatively small introns (mean paradise fish intron length = 566 bp, whereas mean average teleost intron size = 1,214 [22]) (Figure 1B) and shorter intergenic regions [21]. The N90 for our assembly consists of 23 contigs suggesting that most chromosomes are primarily represented by a single contig from the *de novo* assembly, even without any scaffolding performed. Searching the contigs with telomeric sequences revealed “telomere-to-telomere” assemblies for contigs ptg000004l, ptg000010l, ptg000024l, ptg000026l, ptg000028l, and ptg000030l representing chromosomes 3, 8, 9, 15, 17 and 21, respectively (Supplementary Table S2). These contigs have vertebrate telomeres at both ends while the remaining contigs have one or no stretches of telomeric sequence at the end of the contig.

Benchmarking Universal Single-Copy Orthologs (BUSCO) was used to evaluate the completeness of our reference genome assembly with the Actinopterygii_odb10 dataset [23, 24]. The result showed that 98.5 % of the sequence in the reference dataset had a complete ortholog in our genome including 97.3% complete and single-copy genes and 1.2% complete and duplicate genes. Additionally, 1.2 % of the genes were reported as fragmented and 0.3 % of the genes were completely missing.

### Paradise fish genome assembly repeat content characterization

Using repeatmasker [25], we analysed and characterized the repeat content in our reference genome assembly. By using a custom-built repeat prediction library, we identified 32,955,420 bp (6.78 %) in retroelements and 11,076,209 bp (2.8 %) in DNA transposons (Supplementary Table, S3). The retroelements were further categorised into repeat families which were made up of short interspersed nuclear elements (SINEs), long interspersed nuclear elements (LINEs), or long terminal repeats (LTR) (supplementary Table, S3). The LINEs were the most abundant repetitive sequence in the retroelement family at 3.38 % (16,447,763 bp) followed by LTRs, 3.19 % (15,490,642 bp), and SINEs occurred at a lowest frequency (0.21 %) (supplementary Table, S3). In the LINEs sub-family, we identified L2/CR1/Rex as the most abundant repetitive sequence (2.15 %) followed closely with the retroviral (1.73 %) LTR sub-family (Supplementary Table, S3). The proportion of the DNA transposons was estimated to be 11,076,209 bp (2.28 %). Overall, the proportion of retroelements (6.78 %) was much higher in the genome compared to that of DNA transposons (2.28 %).

### Transcriptome assembly and quality assessment

The Trinity transcriptome assembler was used to assemble the Illumina short reads from the RNA-sequencing data [26] into predicted transcripts. The transcriptome assembly consisted of 366,029 contigs in 20,157 loci. The integrity of the transcriptome assembly was evaluated by mapping the Illumina short reads back to the assembled transcriptome using bowtie2 [27]; a 98.4 % overall alignment rate was achieved. The BUSCO analysis confirmed 99.6% completeness with 8.2% single copy orthologs and 91.4% duplicated genes (*i.e*. multiple isoforms). A total of 0.4% of the genes were fragmented and 0.0% were missing completely.

### Genome annotation

We analyzed the predicted genes using OrthoFinder [28] compared to the *Betta splendens* [16], medaka [29] and zebrafish [30] genomes (Figure 2). Our analysis shows that 89.6% of the predicted genes (18,057/20,157) of paradise fish could be assigned to orthogroups (Figure 2B), of which only a very low percentage – 2.5% (511/20,517) – were present in species-specific orthogroups (Figure 2C). A vast majority of the annotated genes (17,546/20,517) had orthologs in at least one of the analyzed species, with 70% (14,067/20,517) having orthologs in all the other species (Figure 2D). The ratio of shared orthogroups also supports the expected phylogeny (Figure 2E).

**Figure 2:**
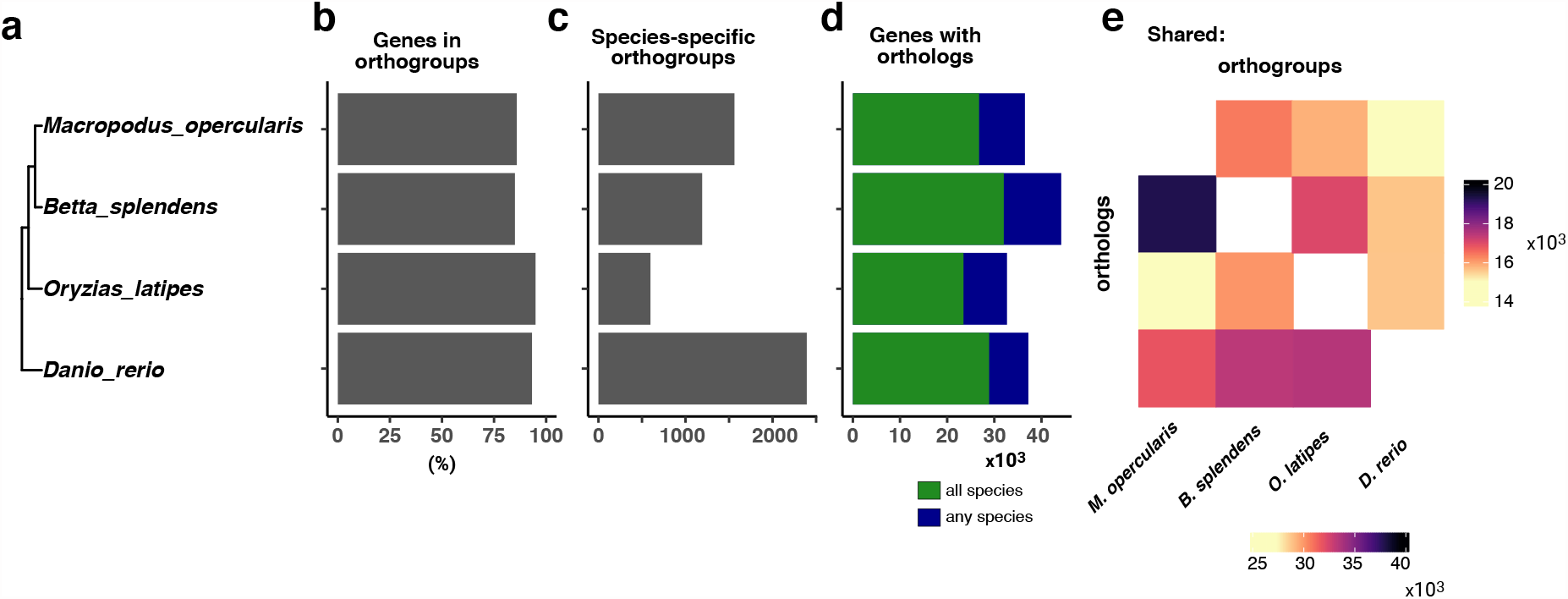
Comparison of predicted proteins across 4 species. A) Evolutionary relationship between paradise fish, Siamese fighting fish, zebrafish, and Japanese medaka. B) The number of genes per species that could be placed in an orthogroup (the set of genes descended from a single gene in the last common ancestor of all the species). This value is close to 90% for all four species. C) The number of orthogroups that are specific to each species. D) The total number of transcripts with orthologs in at least one other species. E) Heat map of the orthogroups for each species pair (top) and orthologs between each species (bottom).

The Vertebrate Genome Project (VGP) [31] had performed short read sequencing on a single paradise fish purchased from a German pet shop (NCBI accession: PRJEB19273), and we captured 9 “wild” samples from the New Territories in Hong Kong. We performed short read sequencing to ≥20X coverage for each fish and used the data to establish the SNP rates within the paradise fish populations. From a single fish (VGP), we identified 2,356,443 variants having a quality score of greater or equal to 30 (Table 1). The transition/transversion rate was 1.35. Our analysis identified a total of 623,414 insertions or deletions ranging from 1 to 60 bps. The rate of SNPs and the indels were 0.5% and 0.1%, respectively.

**Table 1:**
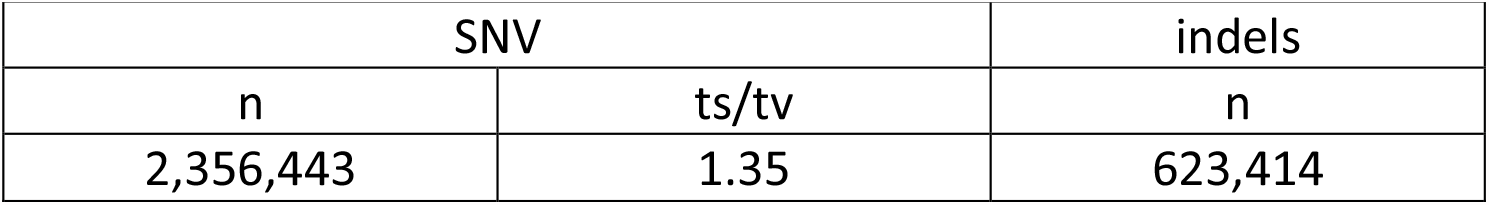
Paradise fish variants call summary statistics from one fish compared to the reference assembly. The single nucleotide variants, SNV, transitions/transversions, ts/tv, and insertions and deletions (indels).

### Cloning and expressional analysis of developmentally important genes

Finally, we selected the (predicted) orthologs of some genes known to be important for early zebrafish development and tested their larval expression by *in situ* hybridization. We successfully cloned fragments of the genes encoding germ plasm determinant Bucky ball (Buc) [32], the dorsally expressed BMP-antagonist Chordin (Chd) [33], the dorsal homeobox transcription factor Goosecoid (Gsc) [34], the axially expressed T-box homeobox transcription factor Tbxta [35, 36], the myogenic differentiation factor MyoD [37], the paired-box transcription factor Pax2a active in the embryonic brain [38] and the eye-fate specifier Retinal homebox 3 (Rx3) [39]. The expression of all these factors (Figure 3) was highly similar to the ones observed in zebrafish, highlighting their conserved role in teleost development.

**Figure 3:**
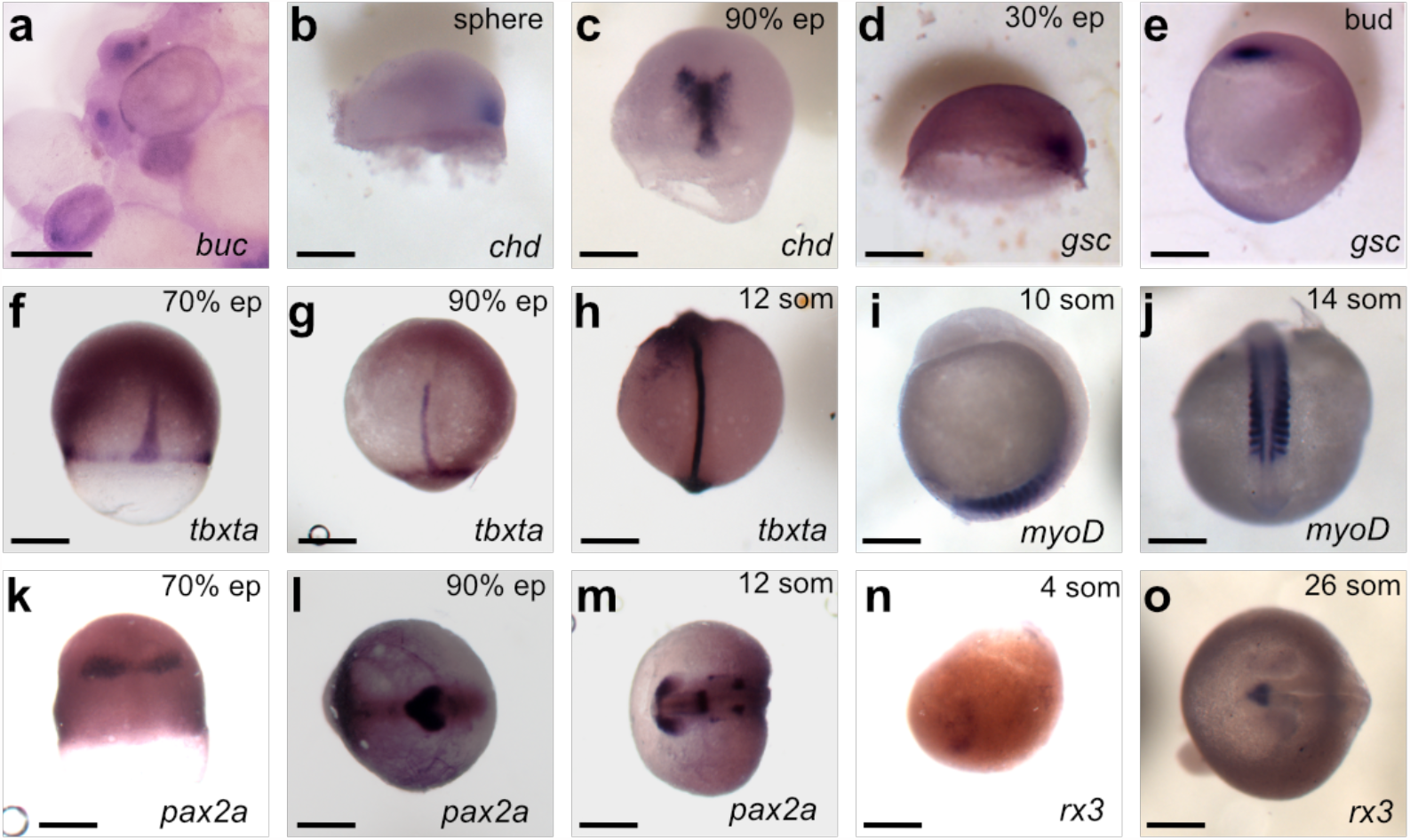
Expression of developmentally important genes in paradise fish embryos visualized by whole-mount *in situ* hybridization. cDNAs of several genes known to regulate early development were isolated from *M. opercularis* and used to make probes for hybridization. A-O) Spatiotemporal expression patterns of all genes are consistent with the zebrafish homologs. Developmental stages (upper right) and gene symbols (lower right) are listed for each panel.

## Conclusions

We report here a high-quality reference genome assembly for *Macropodus opercularis*, commonly known as the paradise fish. Paradise fish have 23 chromosomes and 90% of the total assembly was contained in 23 contigs suggesting that for relatively small genomes a high depth of PacBio HiFi data appears to be sufficient for nearly complete assembly of the genome.

We have generated a detailed transcriptome using RNA sequences from embryonic samples, regenerating fins and adult organs. Over 85.5% of the annotated genes in the *M. opercularis* genome have orthologs at least in one of the three other species (zebrafish, Japanese medaka and *B. splendens*) used in our comparative approach, demonstrating that paradise fish will compliment other model organism research. The *in situ* hybridization results indicate that the spatiotemporal expression patterns of homologous genes are consistent with a conserved role in teleost development. Altogether, our results show that the paradise fish has a small genome with shorter than average intron sizes, as such this could be not only an excellent resource for future neurogenetic studies, but an attractive model organism for genomic research as well.

## Methods

### Animals and husbandry conditions

The founder fish for our colony and the source of the transcriptome samples were purchased from a local pet store. Adult paradise fish were kept in aerated glass aquariums in the animal facility of the Institute of Biology at ELTE Eötvös Loránd University. Husbandry conditions were specified previously [12] Embryos were raised at 28.5°C and staged as described before [40] All experimental procedures were approved by the Hungarian National Food Chain Safety Office (Permit Number: PE/EA/406—7/2020).

### Sample collection, library preparation and sequencing

RNA samples were collected from a mix of embryonic stages (stage 9 – 5 days post fertilization), from caudal tail blastema taken at 3-and 5-days post amputation, from the kidney, heart, brain, ovaries of an adult female, and the brain and testis of an adult male paradise fish, respectively. Total RNA was isolated using TRIzol (Invitrogen, 15596026), following the manufacturer’s protocol. Samples were purified twice with ethanol and eluted in water. Quality and integrity of the samples was tested on an agarose gel, by Nanodrop, and using an Agilent 2100. Ribosomal RNA (rRNA) was removed using the Illumina Ribo-Zero kit and paired-end (PE) libraries were prepared using standard Illumina protocols. Samples were processed on an Illumina NovaSeq PE150 platform, and a total of 218,715,409 PE reads (2x 150 bp) were sequenced, resulting in ∼65 Gbp of raw transcriptomic data.

Genomic DNA samples were isolated from the tail fin of the parental F_0_ male and female paradise fish using the Qiagen DNeasy Blood and Tissue Kit (cat no: 69504). Samples were eluted in TE and sent for library preparation and sequencing. Sample quality-checks were performed using standard agarose gel electrophoresis and with a Qubit 2.0 instrument. For Illumina short-read sequencing a size-selected 150 bp insert DNA library was prepared and processed on the Illumina NovaSeq 5000 platform. Approximately 100 million PE reads (2×150 bp) were sequenced for each parent, resulting in approximately 60X coverage for each genome. For PacBio HiFi long-read single molecule real-time (SMRT) sequencing libraries, genomic DNA was prepared using whole tissue from the F_1_ offspring and the Circulomics Nanobind tissue kit. Sequence libraries were prepared using the PacBio SMRTbell Template Preparation Kit and HiFi sequenced on a Sequel II platform. A total of 4,885,238 reads (average length: 15.5 kbp) resulted in ≈73 Gbp of raw genomic sequence data.

### Genome assembly

All software versions used are listed in Supplementary Table S4. The raw data pre-processing was conducted by doing quality control, adapter trimming, and filtration of the low-quality reads using time_galore!, wrapper around FASTQC and Cutadapt [41]. The genome assembly was generated with the hifiasm genome assembler [18] using the High-Performance Computing facility at the National Institute of Health. For the assembly, 32 cores processing units and 512 Gb of memory was used. The lower and the upper bound binned K-mers was set to 25 and 75, respectively. The estimated haploid genome size used for inferring reads depth was set to 0.5Gbp. The rest of the hifiasm default settings were used to assemble the homozygous genome with the build-in duplication purging parameter set to -l1. The primary assembly Graphical Fragment Assembly (GFA) file was converted to FASTA file format using the awk command.

### Genome annotation

The Trinity assembler [26] was used to create a set of RNA transcripts from the bulk RNA-seq data. To aid in gene prediction, we downloaded the reviewed Swissprot/Uniprot vertebrate proteins (Download date 12/01/2022; entries 97,804 proteins) for homology comparisons in annotation pipelines. Gene prediction was done using the AUGUSTUS [42] and GeneMark-ES [43] softwares as part of the BRAKER pipeline [44] to train the AUGUSTUS parameters. Final annotation using the assembled transcripts and the vertebrate proteins database was done using the MAKER pipeline [45] with the EvidenceModeller [46] tool switched-on to improve gene structure annotation.

### Data for comparative statistics

Sources for the reference data used to create Figure 1B: Refs. [22], [47], [48], [49], [50].

### OrthoFinder analysis

We performed OrthoFinder analysis [28, 51] with default parameters, using predicted peptides (including all alternative splice versions) of the zebrafish genome assembly GRCz11, medaka genome assembly ASM223467v1 and *B. splendens* genome assembly fBetSpl5.3. Sequences were downloaded from the ENSEMBL and NIH/NCBI Assembly homepages, respectively.

### cDNA synthesis, primers, antisense probe synthesis and *in situ* hybridization

cDNA was synthetized using Invitrogen™ Second Strand cDNA Synthesis Kit (A48570) from the total RNA sample mixtures. Selected genes were amplified with DreamTaq™ DNA polymerase (Thermo Scientific, A48571). Primer sequences for each gene were designed based on the *in silico* predictions. PCR amplified fragments were cloned into the pGEMT-Easy vector. For probe synthesis, plasmids were digested and transcribed with either T7 or SP6 RNA polymerases (Thermo Scientific, EP0111 and EP0131, respectively). RNA probes were labeled using digoxigenin labelled dNTPs (Sigma-Aldrich, 11277073910).

We adapted a well-established zebrafish *in situ* hybridization protocol [52], with some specific modifications. Most of these modifications are related to the presence of an oil droplet within paradise fish embryos that keeps them afloat in the bubble nest during early phases of development. Early embryos (before the end of the epiboly) were fixed using Bouin solution, whereas for later stage embryos we used 4% formaldehyde. After 24 hours at 4°C the fixative was washed away, and the chorion was removed using Proteinase K digestion. Early and later stage embryos were stored at -20°C in 75% ethanol and 100% methanol, respectively. The embryos were quite fragile so they were kept in specialized baskets during probe hybridization.

### Variant calling

The Illumina short read files (accession number ERR3332352) were downloaded from the Vertebrate Genome Project (VGP) database (https://vertebrategenomesproject.org/). Trim galore version 0.6.10 (https://github.com/FelixKrueger/TrimGalore), a wrapper around cutadapt and fastqc was then used to trim the illumina adapter sequences and to discard reads less than 25bps. DRAGMAP version 1.3.0 (https://github.com/Illumina/DRAGMAP) was used to map the reads to the reference genome. The resultant sequence alignment map (SAM) file was then converted to binary alignment map (BAM), sorted, and indexed using samtools. The Picard was then used to add the read groups information in BAM file. The genome analysis tool kit (GATK) was then used in calling the variants by turning on the dragen mode. The bamtools stats and the plot-vcfstats was used in the downstream analysis and visualization of the genomic variants in the variant call file (VCF).

### Availability of data and materials

The assembly and all DNA and RNA raw reads have been deposited in the NCBI under the BioProject study accession PRJNA824432. The sequence of amplified cDNA fragments used in the *in-situ* hybridization experiments are available at Zenodo [53].

## Authors’ contributions

Conceptualization: ÁM, LO, SB, MV.

Data curation: JO, MV, SB.

Funding acquisition: ÁM, SB, MV.

Investigation: EF, JO, NS, KS, DC, AR, SK, AR.

Methodology: EF, JO, NS, DC, ISz, LO, MV, SK, AR.

Project administration and supervision: SB, MV.

Writing – original and revised text: EF, JO, SB, MV.

## Acknowledgements

The authors thank Lars Martin Jakt and his team for early access to their unpublished data. This work utilized the computational resources of the NIH HPC Biowulf cluster (http://hpc.nih.gov). We would like to thank Adam Phillippy and Brandon Pickett for helpful discussions.

